# Bidirectional modulation of aging-associated cellular phenotypes by mitochondrial genome replacement

**DOI:** 10.64898/2026.06.26.734763

**Authors:** Fumiya Teruyama, Toshihiko Taya, Akira Shikuma, Satoaki Matoba, Kentaro Numajiri, Sayako Umetani, Yutong Chen, Amnon Peled, Eithan Galun, Nimer Assy, Aviad Zick, Silvia Noiman, Dayana Michel, Dori Pelled, Taro Inaba, Kenji Chamoto, Satoshi Gojo

**Affiliations:** Department of Regenerative Medicine, Graduate School of Medicine, Kyoto Prefectural University of Medicine, Kyoto, Japan; Department of Cardiovascular Medicine, Graduate School of Medicine, Kyoto Prefectural University of Medicine, Kyoto, Japan; Tokyo New Drug Research Laboratories, Kowa Company, Ltd., Tokyo, Japan; Bio Science & Engineering Laboratories, FUJI FILM Corporation, Tokyo, Japan; Division of Immunology and Genomic Medicine, Center for Cancer Immunotherapy and Immunobiology, Graduate School of Medicine, Kyoto University; Goldyne Savad Institute of Gene Therapy& Sharett Institute of Oncology, Hadassah Hebrew University Hospital, Jerusalem, Israel; Internal Medicine Department, Galilee Medical Center Hospital, Nahariya-Cabri, Israel; Sharett Institute of Oncology, Hadassah Medical Center and the Faculty of Medicine, The Hebrew University of Jerusalem, Jerusalem, Israel; IMEL Biotherapeutics, Inc, MA, USA; IMEL Japan, Inc, Tokyo, Japan; Division of Cancer Immune Regulation, Center for Cancer Immunotherapy and Immunobiology, Graduate School of Medicine, Kyoto University

## Abstract

Mitochondrial dysfunction is a hallmark of cellular aging, but whether age-associated cellular decline can be functionally reversed remains unclear. Here, we applied mitochondrial genome replacement to replicative senescent fibroblasts and aged T cells derived from mice and humans. In senescent fibroblasts, replacement with mitochondria from young cells extended proliferative lifespan, whereas replacement with aged mitochondria accelerated proliferative decline, indicating bidirectional modulation of aging-associated phenotypes. In aged mouse T cells, mitochondrial genome replacement restored proliferative capacity and significantly enhanced antitumor activity following adoptive transfer into tumor-bearing mice. Similarly, mitochondrial genome replacement in aged human T cells enhanced cytokine production and shifted the transcriptomic programs toward a more youthful state. Collectively, these findings identify mitochondrial genetic integrity as a functional regulator of aging-associated cellular states and support the emerging view that mitochondria actively influence cellular aging trajectories.

## Introduction

Aging is accompanied by progressive functional decline across multiple tissues and organ systems. Among the hallmarks of aging, mitochondrial dysfunction has emerged as a central driver of age-associated cellular deterioration (Lopez-Otin *et al*, 2023). Mitochondria regulate numerous aging-related processes, including metabolic homeostasis (Amorim *et al*, 2022), reactive oxygen species (ROS) production (Sies & Jones, 2020), inflammatory signaling (Riley & Tait, 2020), and quality control pathways such as mitophagy (Uoselis *et al*, 2023). Accumulation of mitochondrial dysfunction contributes to chronic inflammation, impaired tissue maintenance, and progression of age-associated diseases (Suomalainen & Nunnari, 2024).

Replicative senescence in cultured fibroblasts has long served as an experimental model of aging-associated cellular dysfunction, despite known differences between in vitro senescence and organismal aging (Maier & Westendorp, 2009; Rubin, 2002). In parallel, T-cell aging has emerged as a major determinant of organismal aging and age-associated immune dysfunction (Mittelbrunn & Kroemer, 2021). Aged T cells exhibit impaired proliferation, reduced antitumor activity, and chronic inflammatory phenotypes collectively termed immunosenescence and inflammaging (Lee *et al*, 2021; Liu *et al*, 2023). Recent studies demonstrated that transfer of dysfunctional T cells or T cells harboring defective mitochondria induces systemic aging phenotypes, whereas young immune cells can partially reverse aging-associated dysfunction (Desdin-Mico *et al*, 2020; Yousefzadeh *et al*, 2021). Mitochondrial dysfunction is therefore considered a central contributor to T-cell and systemic aging (Escrig-Larena *et al*, 2023).

Until the discovery that mitochondria could move between cells through tunneling nanotubes (Rustom *et al*, 2004), mitochondria were generally regarded as static intracellular organelles dedicated primarily to energy metabolism. Although horizontal gene transfer had long been recognized in prokaryotes (Watanabe & Fukasawa, 1961), transfer of mitochondrial genomes between eukaryotic cells was not considered biologically relevant. Subsequent studies demonstrating that intercellular mitochondrial transfer could restore cellular phenotypes in vitro (Spees *et al*, 2006) fundamentally expanded the concept of mitochondrial biology. Mitochondria are increasingly recognized as dynamic organelles capable of intercellular transfer. Mitochondrial transfer has been observed in multiple biological contexts, including stromal-to-epithelial and astrocyte-to-neuron communication (Hayakawa *et al*, 2016; Islam *et al*, 2012). In parallel, isolated mitochondria can be internalized through macropinocytosis (Kitani *et al*, 2014), enabling experimental mitochondrial transplantation approaches. We previously developed a mitochondrial genome replacement platform in which recipient cells are transiently rendered mtDNA-reduced (ρ(-)) cells prior to co-culture with isolated donor mitochondria (Maeda *et al*, 2021). Previous studies demonstrated that donor mitochondrial nucleoids released from internalized mitochondria can migrate into recipient mitochondrial matrices following mitochondrial uptake, resulting in stable replacement of endogenous mitochondrial genomes (Maeda *et al*., 2021). This process restored mitochondrial function in cells carrying pathogenic mtDNA mutations.

Given the central role of mitochondrial dysfunction in T-cell aging, we hypothesized that restoration of mitochondrial genetic integrity through mitochondrial genome replacement may partially reverse aging-associated cellular dysfunction. In this study, we establish mitochondrial genome replacement in primary mouse and human T cells and investigate its effects on proliferation, antitumor activity, mitochondrial function, and transcriptomic remodeling in aged T cells.

## Results

### Generation of mitochondrial genome-replaced fibroblasts with donor-derived mtDNA dominance

To investigate the contribution of mitochondrial genetic integrity to aging-associated cellular phenotypes, recipient fibroblasts were transiently rendered mtDNA-reduced (ρ(-)) through expression of a mitochondrially targeted endonuclease (MTS-Xbal) and subsequently co-cultured with isolated donor mitochondria, generating mitochondrial genome-replaced cells (MirCs) as previously described (Maeda *et al*., 2021) (Fig. 1A). To validate mitochondrial genome replacement in fibroblasts, we used two primary human fibroblast lines, TIG-1 and NHDF, which carry distinct mtDNA SNPs at position 16145 (Fig. 1B and EV1A). Following co-culture of mtDNA-reduced NHDF cells with isolated TIG-1 mitochondria, sequencing analysis and TaqMan SNP assays demonstrated that donor-derived mtDNA accounted for >90% of total mitochondrial genomes in MirCs (Fig. 1B and EV1A,B).

**Figure 1.**
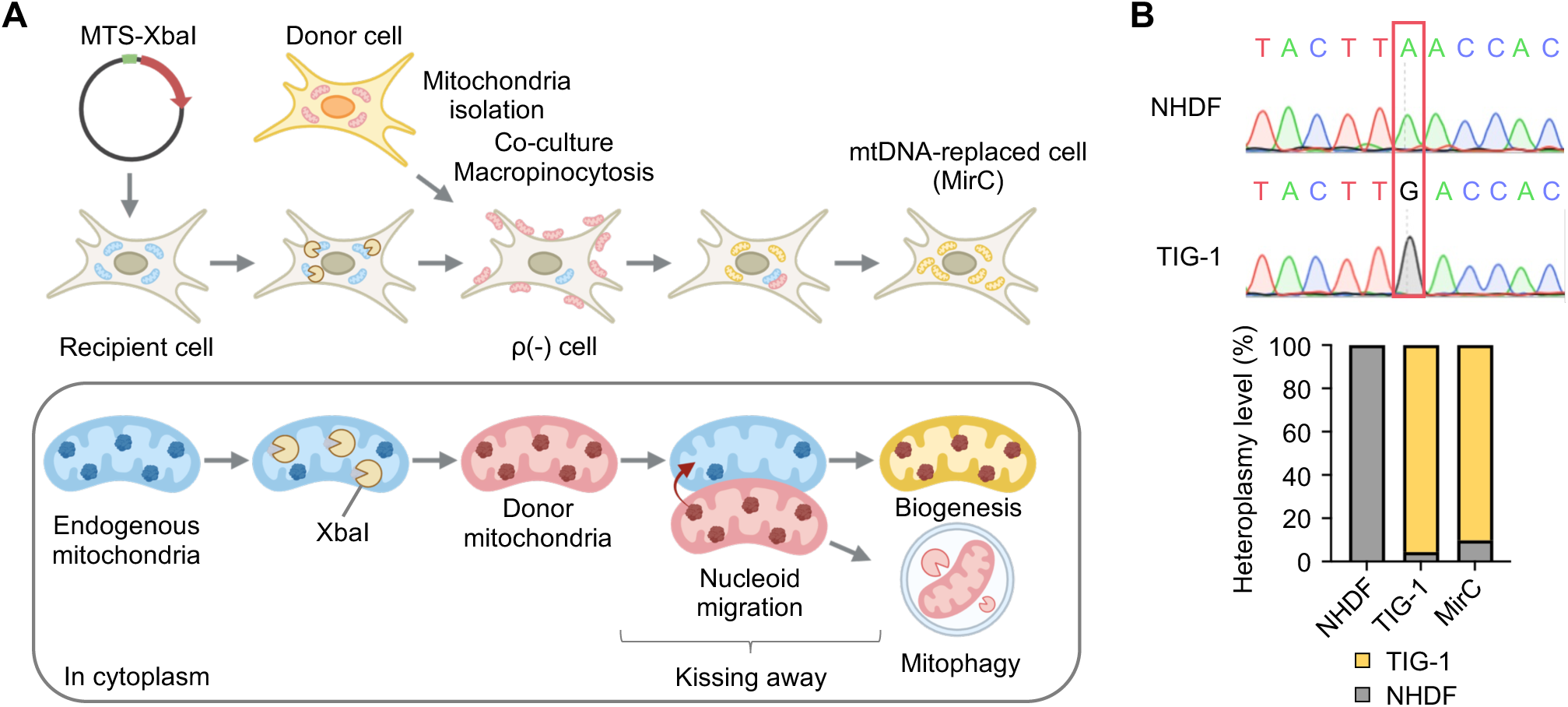
Generation and validation of mitochondrial genome-replaced fibroblasts. (A) Schematic overview of mitochondrial genome replacement in fibroblasts. Recipient cells were converted into mtDNA-reduced ρ(-) cells through expression of a mitochondrially targeted endonuclease and subsequently co-cultured with isolated donor mitochondria to generate mitochondrial genome-replaced cells (MirCs). The upper panel illustrates the experimental workflow, whereas the lower panel depicts the proposed intracellular process based on previous observations of donor nucleoid migration into recipient mitochondrial matrices. Created with BioRender.com. (B) Validation of donor mitochondrial genome incorporation in MirCs. Representative mtDNA SNP chromatograms from NHDF and TIG-1 cells and heteroplasmy analysis of MirCs determined by TaqMan SNP assays targeting donor-specific mtDNA variants.

### Mitochondrial genome replacement bidirectionally modulates aging-associated phenotypes in fibroblasts

To examine whether mitochondrial genome replacement influences aging-associated cellular phenotypes, we generated MirCs using normal human dermal fibroblasts (NHDFs) at different stages of replicative aging. Cells at population doubling levels (PDLs) of 10–20 were defined as young, whereas cells at PDL ≥40 were defined as aged. Aged NHDFs receiving mitochondrial genomes from young donor cells (NHDF^Young/Aged^) and young NHDFs receiving mitochondrial genomes from aged donor cells (NHDF^Aged/Young^) were generated (Fig. EV1C) and continuously cultured until growth arrest. Replacement of aged mitochondrial genomes with young donor mitochondrial genomes (Young/Aged) extended proliferative lifespan by approximately 10 additional population doublings compared with control aged NHDFs, which underwent growth arrest at approximately PDL 55 (Fig. 2A). In contrast, replacement of young mitochondrial genomes with aged donor mitochondrial genomes (Aged/Young) reduced proliferative lifespan by approximately 10 population doublings (Fig. 2A). These findings indicate that mitochondrial genome replacement bidirectionally modulates replicative aging-associated phenotypes in fibroblasts. To evaluate metabolic alterations following mitochondrial genome replacement, oxygen consumption rate (OCR) was measured using substrate–uncoupler–inhibitor titration assays. NHDF^Young/Aged^ cells cultured for one month after mitochondrial genome replacement exhibited significant increases in routine respiration, maximal respiration, and ATP production, approaching levels observed in young NHDFs (Fig. 2B,C and EV1D). We next assessed senescence-associated β-galactosidase activity by flow cytometry. Elevated β-galactosidase activity observed in aged NHDFs was significantly reduced in NHDF^Young/Aged^ cells (Fig. 2D). Together, these findings suggest that mitochondrial genome replacement partially reverses aging-associated functional and metabolic phenotypes in fibroblasts.

**Figure 2.**
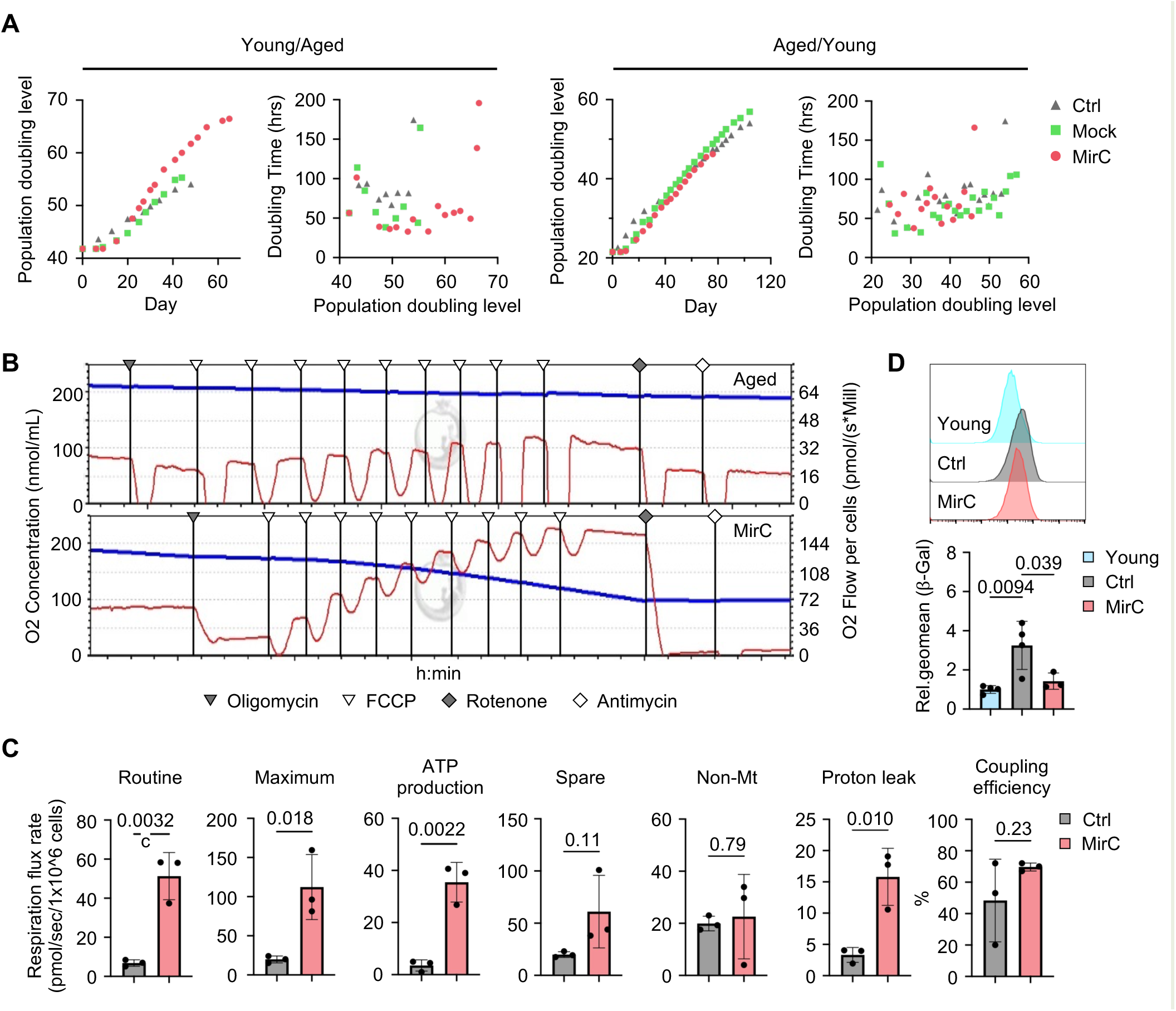
Bidirectional modulation of aging-associated phenotypes by mitochondrial genome replacement in fibroblasts. (A) Cumulative population doublings and doubling time during continuous culture of mitochondrial genome-replaced fibroblasts (MirCs). Left, aged fibroblasts receiving mitochondrial genomes from young donor cells (Young/Aged); right, young fibroblasts receiving mitochondrial genomes from aged donor cells (Aged/Young). Doubling time is plotted against population doubling level (PDL). N = 1. (B) Representative oxygen consumption traces measured by substrate–uncoupler–inhibitor titration using an O2k-FluoRespirometer in MirCs generated by replacement of aged mitochondrial genomes with young donor mitochondrial genomes. Blue line indicates chamber oxygen concentration (left axis), and red line indicates oxygen flux per cell (right axis). Representative traces from three independent experiments are shown. (C) Quantification of respiratory parameters derived from the OCR analysis shown in (B), including routine respiration, maximal respiration, ATP-linked respiration, spare respiratory capacity, non-mitochondrial respiration, proton leak, and coupling efficiency (n = 3). (D) Senescence-associated β-galactosidase staining in MirCs generated by replacement of aged mitochondrial genomes with young donor mitochondrial genomes (n = 3–4). Data in (C) and (D) are presented as mean ± SD. P values were determined using Student’s t-test (C) or one-way ANOVA followed by Tukey’s multiple-comparison test (D).

### Stable mitochondrial genome replacement in primary mouse and human T cells

To extend mitochondrial genome replacement to immune cells, we established a protocol to generate mitochondrial genome-replaced T (MirT) cells from primary mouse and human CD3^+^ T cells. Because prolonged expression of mitochondrially targeted endonucleases may impair cellular function, we employed transient mRNA-based delivery of mitochondrially targeted XbaI endonuclease. In mouse CD3^+^ T cells, electroporation conditions were optimized to achieve near-complete transfection efficiency, as confirmed by mitochondrial-targeting sequence (MTS)-EGFP expression (Fig. 3A). EGFP expression progressively declined and became minimal by day 4 after transfection (Fig. 3B). mtDNA copy number analysis demonstrated a 60–70% reduction in endogenous mitochondrial genomes following MTS-XbaI mRNA transfection (Fig. 3C). Donor mitochondria were subsequently added on day 5 (Fig. EV2A), when endonuclease expression was minimal. To assess mitochondrial genome replacement, recipient T cells from NZB mice and donor mitochondria from B6 mice carrying distinct mtDNA SNPs were used (Fig. EV2B,C). TaqMan SNP assays demonstrated donor-derived mtDNA incorporation reaching approximately 80% at the population level in MirT cells (Fig. EV2D and 3D). Single-cell droplet digital PCR (ddPCR) analysis further revealed that approximately 60% of MirT cells exhibited donor-derived homoplasmy, whereas only a small fraction of T cells retained mixed mitochondrial genotypes (heteroplasmy) (Fig. 3E). We next adapted this protocol to human CD3^+^ T cells using a clinically applicable electroporation platform. Similar to mouse T cells, expression of the introduced endonuclease progressively declined after transfection (Fig. 3F), accompanied by a 60–70% reduction in mtDNA copy number (Fig. 3G). Using donor mitochondria isolated from EPC100 mesenchymal cells carrying distinct mtDNA SNPs (Fig. EV3A-C), TaqMan SNP assays demonstrated donor derived heteroplasmy shifts of 60–70% in human MirT cells (Fig. 3H). Single-cell ddPCR analysis further confirmed donor genotype dominance in the majority of human MirT cells (Fig. 3I). Finally, to minimize inflammatory responses associated with mRNA delivery, pseudouridine-modified mRNA was evaluated. Pseudouridine modification significantly reduced inflammatory cytokine induction following MirT cells generation (Fig. 3J), and was therefore used in subsequent experiments.

**Figure 3.**
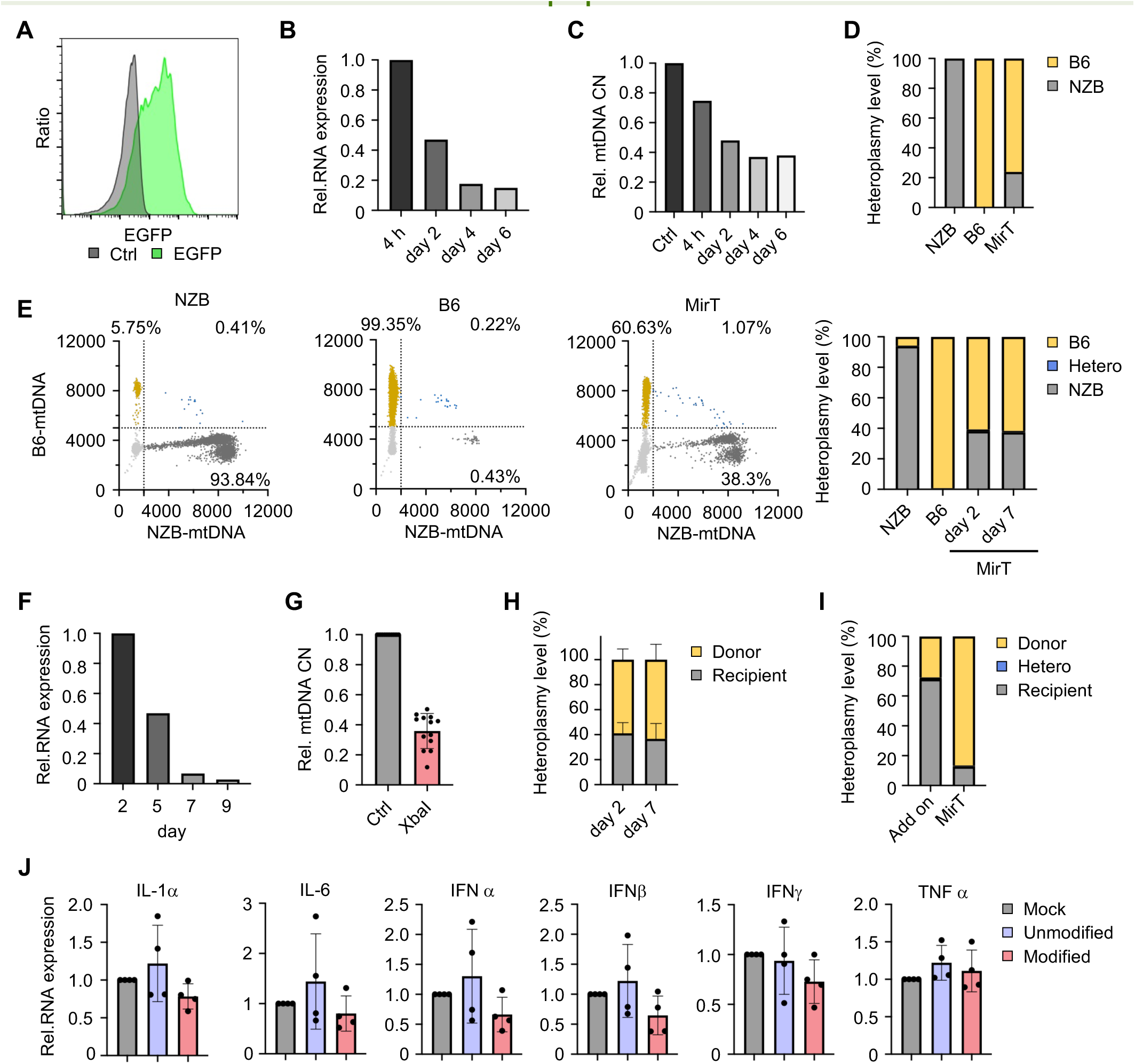
Establishment of mitochondrial genome replacement in primary mouse and human T cells. (A) Representative flow cytometric analysis of EGFP expression 48 h after transfection of mouse CD3+ T cells with MTS-EGFP mRNA. (B) Time course of EGFP expression following MTS-EGFP mRNA transfection in mouse CD3+ T cells. (C) Time course of mtDNA copy number following transfection with MTS-XbaI mRNA in mouse CD3+ T cells. (D) Mitochondrial heteroplasmy in mitochondrial genome-replaced T cells (MirT cells) on day 7, determined by TaqMan SNP assays. (E) Single-cell mitochondrial genotype analysis during MirT cells generation using sc-ddPCR. Representative plots from day 7 are shown. Bar graphs indicate heteroplasmy quantification on days 7 and 12. (F) Time course of EGFP expression following MTS-EGFP mRNA transfection in human CD3+ T cells. (G) mtDNA copy number in human CD3+ T cells 3 days after transfection with MTS-XbaI mRNA. n = 12 independent experiments. (H) Mitochondrial heteroplasmy in human MirT cells on days 7 and 12, determined by TaqMan SNP assays. (I) Single-cell mitochondrial genotype analysis of human MirT cells on day 7 using sc-ddPCR. For comparison, an “Add on” control group was included, in which cells underwent co-culture without prior mitochondrial DNA reduction. (J) Expression of inflammatory cytokines 3 days after pseudouridine-modified mRNA transfection (n = 4). Data in (G), (H), and (J) are presented as mean ± SD.

### Restoration of antitumor function in aged T cells by mitochondrial genome replacement

Having established mitochondrial genome replacement in young T cells, we next examined whether this approach could be extended to aged T cells derived from mice older than 18 months. In contrast to young T cells, aged T cells exhibited markedly reduced survival and transfection efficiency during prolonged culture and electroporation procedures. Unless otherwise specified, “MirT cells” in the following experiments refers to replacement of aged mitochondria with mitochondria derived from young cells (MirT cells^Young/Aged^). To improve gene delivery into aged T cells, the duration of IL-7 pre-stimulation was optimized (Fig. EV4A). Using this protocol (Fig. EV4B), transfection of MTS-XbaI mRNA induced a 60–70% reduction in mtDNA copy number (CN), comparable to that observed in young T cells (Fig. EV4C). Whereas B6 and NZB mice were used to establish mitochondrial genotype replacement in vitro, the subsequent animal studies employed a B6–C3H strain combination. To enable quantification of donor-derived heteroplasmy shifts, strain-specific mtDNA SNPs were identified and TaqMan qPCR assays were developed to distinguish B6 and C3H mitochondrial genomes (Fig. EV4D,E). TaqMan SNP assays demonstrated donor-derived heteroplasmy shifts of approximately 70–80% in MirT cells generated from aged T cells (Fig. 4A). To examine the stability of donor-derived mitochondrial genomes, heteroplasmy levels were monitored over time and compared with cells exposed to isolated mitochondria without prior mtDNA reduction (“Add-on” group) (Fig. 4B). MirT cells maintained substantially higher heteroplasmy levels than the Add-on group throughout the culture period without significant decline over time, indicating stable maintenance of donor-derived mitochondrial genomes. Across independent experiments, heteroplasmy efficiency strongly correlated with the extent of endogenous mtDNA reduction (Fig. 4C), suggesting that efficient depletion of endogenous mitochondrial genomes facilitates donor mtDNA incorporation. We next examined whether mitochondrial genome replacement could restore antitumor activity in aged T cells: MC38-OVA-bearing mice were treated by adoptive cell transfer (ACT) of OT-1 T cells (Fig. 4D). Oxygen consumption rate (OCR) and extracellular acidification rate (ECAR) analyses were performed in young T cells, aged T cells, and MirT cells cultured under IL-7 maintenance conditions for 5 days according to the MirT cells generation protocol (Fig. EV4B). Even under homeostatic IL-7 conditions, aged T cells exhibited elevated oxygen consumption compared with young T cells (Fig. 4E). In contrast, MirT cells showed normalization of SRC, proton leak, and ATP production toward levels observed in young T cells (Fig. 4E-G). These data demonstrated that MirT cells shifted the metabolic state of aged T cells toward a profile resembling that of young T cells.

**Figure 4.**
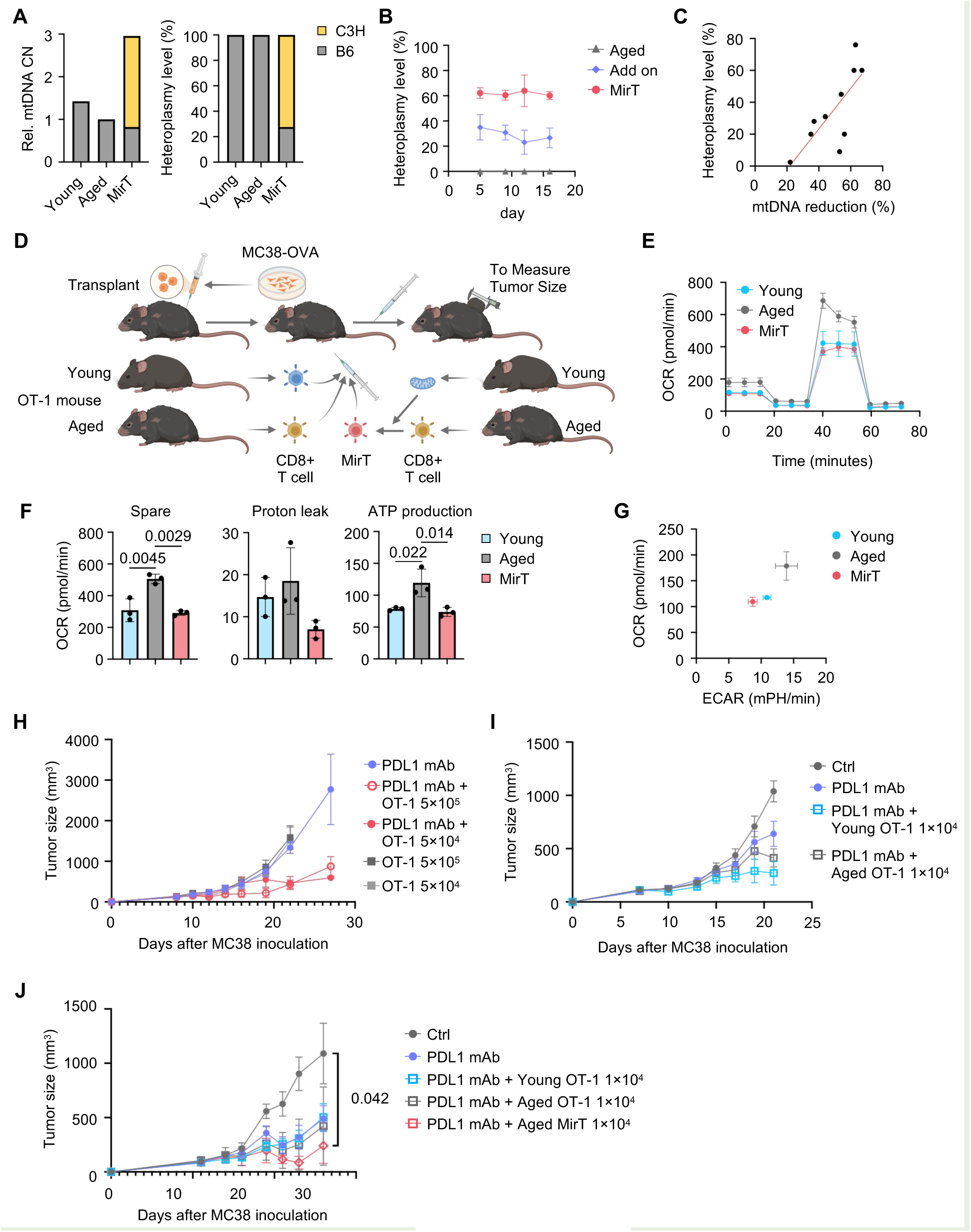
Mitochondrial genome replacement restores antitumor function in aged mouse T cells. (A) mtDNA heteroplasmy in MirT cells on day 5, determined by TaqMan SNP assays. (B) Time course of mitochondrial heteroplasmy following MirT cells generation (n = 3). (C) Correlation between reduction of endogenous mtDNA copy number and donor mtDNA incorporation efficiency. (D) Schematic overview of the adoptive cell transfer (ACT) experiments using OT-1 T cells in MC38 tumor-bearing mice treated with anti-PD-L1 antibody. Created with BioRender.com. (E) Representative OCR traces of Young, Aged, and MirT cells-treated T cells cultured under IL-7 maintenance conditions for 5 days following the MirT cells generation protocol. (F) Quantification of spare respiratory capacity (SRC), proton leak, ATP production (G) OCR/ECAR metabolic profiles derived from the OCR and ECAR analyses shown in panel F. (H) Determination of the number of aged OT-1 T cells required to suppress tumor growth following ACT (n = 3–4). (I) Tumor growth following ACT using young or aged OT-1 T cells in combination with anti-PD-L1 antibody (n = 3–4). (J) Comparison of antitumor activity between young OT-1 T cells, aged OT-1 T cells, and MirT cells following ACT in MC38 tumor-bearing mice treated with anti-PD-L1 antibody. Data are presented as mean ± SD in (E) and mean ± SEM in (H–K). P values were determined using one-way ANOVA followed by Tukey’s multiple-comparison test (F) or two-way mixed-design ANOVA followed by Tukey’s multiple-comparison test (J).

We next evaluated in vivo the effects of ACT and anti-PD-L1 treatment individually and in combination. A tumor model was established in which combined OT-1 cell transfer and anti-PD-L1 treatment produced a more evident antitumor effect than either treatment alone (Fig. 4H). Using this sensitized ACT/anti-PD-L1 combination model, young T cells suppressed tumor growth efficiently, while aged T cells exhibited markedly impaired antitumor activity (Fig. 4I). Donor-derived heteroplasmy levels reached approximately 70% prior to transplantation, as determined by TaqMan SNP analysis (Fig. EV4F). Notably, MirT cells restored antitumor activity to levels comparable to those of young T cells (Fig. 4J), suggesting that mitochondrial genome replacement partially reverses aging-associated T-cell dysfunction in vivo.

### LNP-assisted mitochondrial genome replacement enhances proliferative fitness of aged mouse T cells

Generation of MirT cells from aged T cells using electroporation resulted in substantial cell loss, with viability decreasing below 40% by 2 days after mitochondrial co-culture (Fig. 5A). To improve cell survival, we replaced electroporation with lipid nanoparticle (LNP)-mediated mRNA delivery, using a newly established protocol for LNP-assisted MirT cell generation (Fig. 5B). LNP-mediated delivery reduced mtDNA CN comparable to electroporation-based protocols on Day 5 (Fig. 5C).

**Figure 5.**
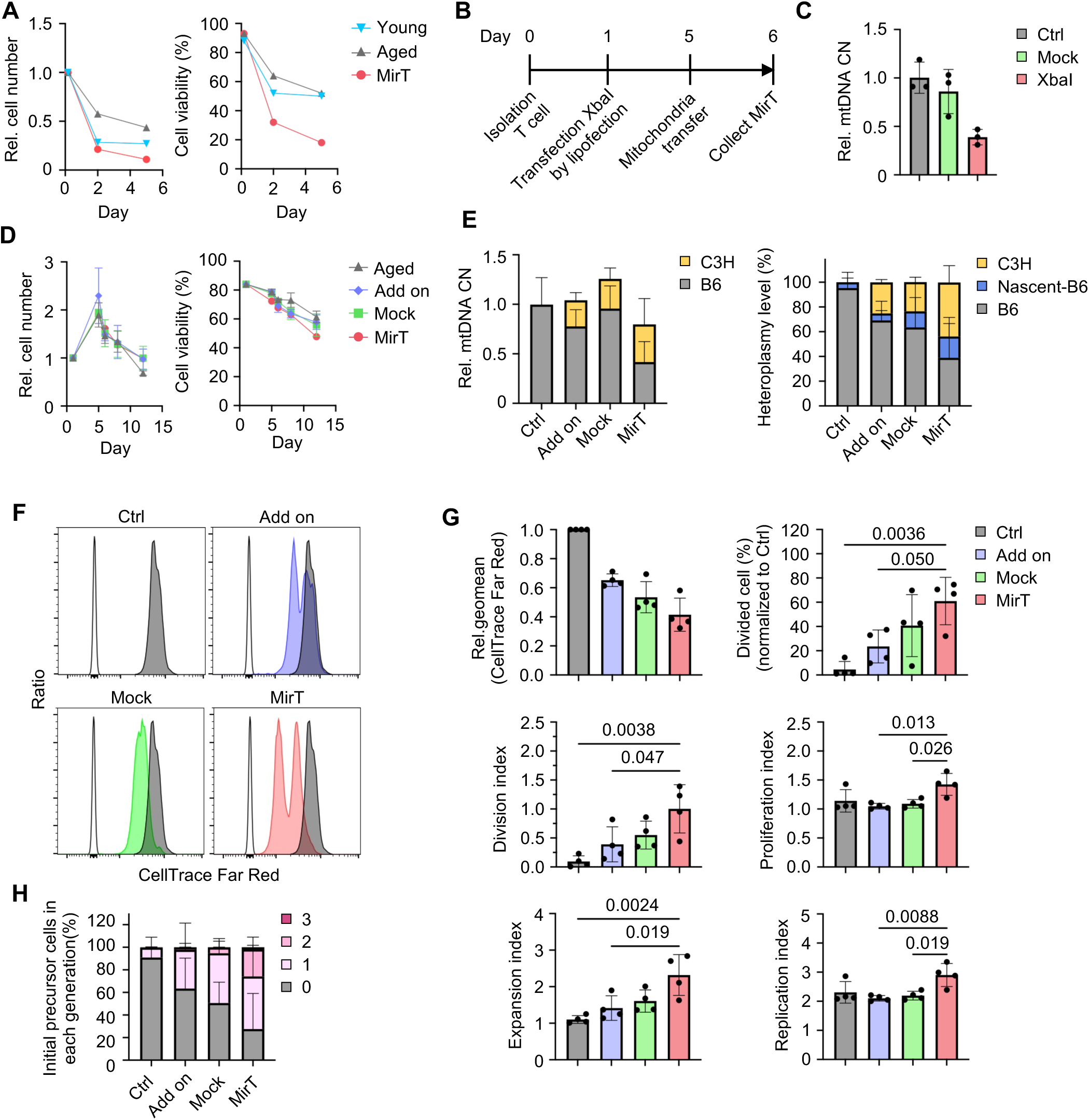
LNP-assisted mitochondrial genome replacement enhances proliferative fitness of aged mouse T cells. (A) Time-course changes in cell number and viability during generation of mitochondrial genome-replaced T cells (MirT cells) from aged mouse T cells. (B) Experimental protocol for LNP-assisted generation of MirT cells from aged mouse T cells. (C) mtDNA copy number following MTS-XbaI mRNA transfection compared with untreated (Ctrl) and MTS-EGFP mRNA-transfected (Mock) control cells. (D) Time-course changes in cell number and viability during LNP-assisted MirT cells generation (n = 3). (E) mtDNA heteroplasmy in MirT cells generated from aged mouse T cells, determined by TaqMan SNP assays (n = 3). (F) Representative CellTrace Far Red proliferation profiles of aged mouse T cells following MirT cells generation (n = 4 independent experiments). (G) Quantification of proliferation parameters derived from CellTrace Far Red assays, including geometric mean fluorescence intensity, percentage of divided cells, division index, proliferation index, expansion index, and replication index (n = 4). (H) Distribution of precursor cells according to generation number based on CellTrace Far Red proliferation analysis. Data in (C), (D), and (G) are presented as mean ± SD. P values were determined using one-way ANOVA followed by Tukey’s multiple-comparison test.

LNP-assisted MirT cells generation markedly improved cell recovery and viability compared with electroporation-based protocols (Fig. 5D). TaqMan SNP assays demonstrated donor-derived heteroplasmy levels of approximately 50% in MirT cells on Day 6, substantially higher than those observed in mock or add on controls (Fig. 5E). Furthermore, analysis of endogenous mtDNA copy number revealed a significant increase in newly synthesized (nascent) mtDNA between Days 5 and 6 in the MirT group. Additional analyses demonstrated that MirT cells did not exhibit overt increases in ROS production, inflammatory cytokine expression, senescence-associated markers, or exhaustion-associated phenotypes during short-term culture, supporting the conclusion that the observed functional effects were not attributable to procedure-induced cellular stress or perturbation of T-cell state (Appendix Fig. S1A-I). We next examined whether mitochondrial genome replacement restores proliferative competence in aged T cells. CellTrace Far Red proliferation assays demonstrated markedly enhanced proliferation in MirT cells compared with control, mock, and add-on groups following CD3/CD28 stimulation (Fig. 5F). MirT cells exhibited enhanced proliferative capacity, as evidenced by higher division, proliferation, expansion, and replication indices (Fig. 5G). Consistent with this, cells undergoing multiple rounds of division (>2 divisions) were predominantly observed in the MirT group (Fig. 5H). Together, these findings indicate that LNP-assisted mitochondrial genome replacement restores proliferative fitness in aged T cells without inducing overt stress-associated phenotypes.

### Functional restoration of aged human T cells following mitochondrial genome replacement

Functional restoration observed in aged mouse MirT cells prompted us to investigate whether mitochondrial genome replacement similarly improves functional properties of aged human T cells. Two independent cohorts were analyzed: healthy elderly donors and elderly cancer patients receiving mitochondria isolated from young donor lymphocytes. In the healthy elderly cohort (mean age, 80 years; Appendix Table S1), MirT cells exhibited donor-derived heteroplasmy levels exceeding 90%, as determined by whole mitochondrial genome sequencing (Fig. 6A,B). Following CD3/CD28 stimulation, MirT cells exhibited significantly increased cytokine production, including IL-2 and TNF-related cytokines, compared with control aged T cells (Fig. 6C). Analysis of mitochondrial membrane potential using TMRM demonstrated an increased proportion of T cells with low mitochondrial membrane potential in MirT cells, whereas the high-TMRM population remained largely unchanged (Fig. 6D). These findings are consistent with remodeling toward a less differentiated metabolic state and align with the restoration of metabolic flexibility observed in MirT cells in mice (Fig. 4E-G). We next examined MirT cells generated from T cells of elderly cancer patients (Appendix Table S2). MirT cells exhibited increased mtDNA content, as determined by quantification of the mitochondrial 12S rRNA gene (Fig. 6E). Similar to healthy elderly donors, MirT cells exhibited donor-derived heteroplasmy levels exceeding 90% (Fig. 6F). MirT cells derived from elderly cancer patients similarly demonstrated increased low-TMRM populations together with increased mitochondrial mass (Fig. 6G,H). Under unstimulated conditions, MirT cells exhibited reduced basal cytokine expression compared with original aged T cells (Fig. 6I). In contrast, CD3/CD28 stimulation induced markedly increased IL-2 production in MirT cells (Fig. 6J). Together, these findings indicate that mitochondrial genome replacement partially restores functional responsiveness in aged human T cells.

**Figure 6.**
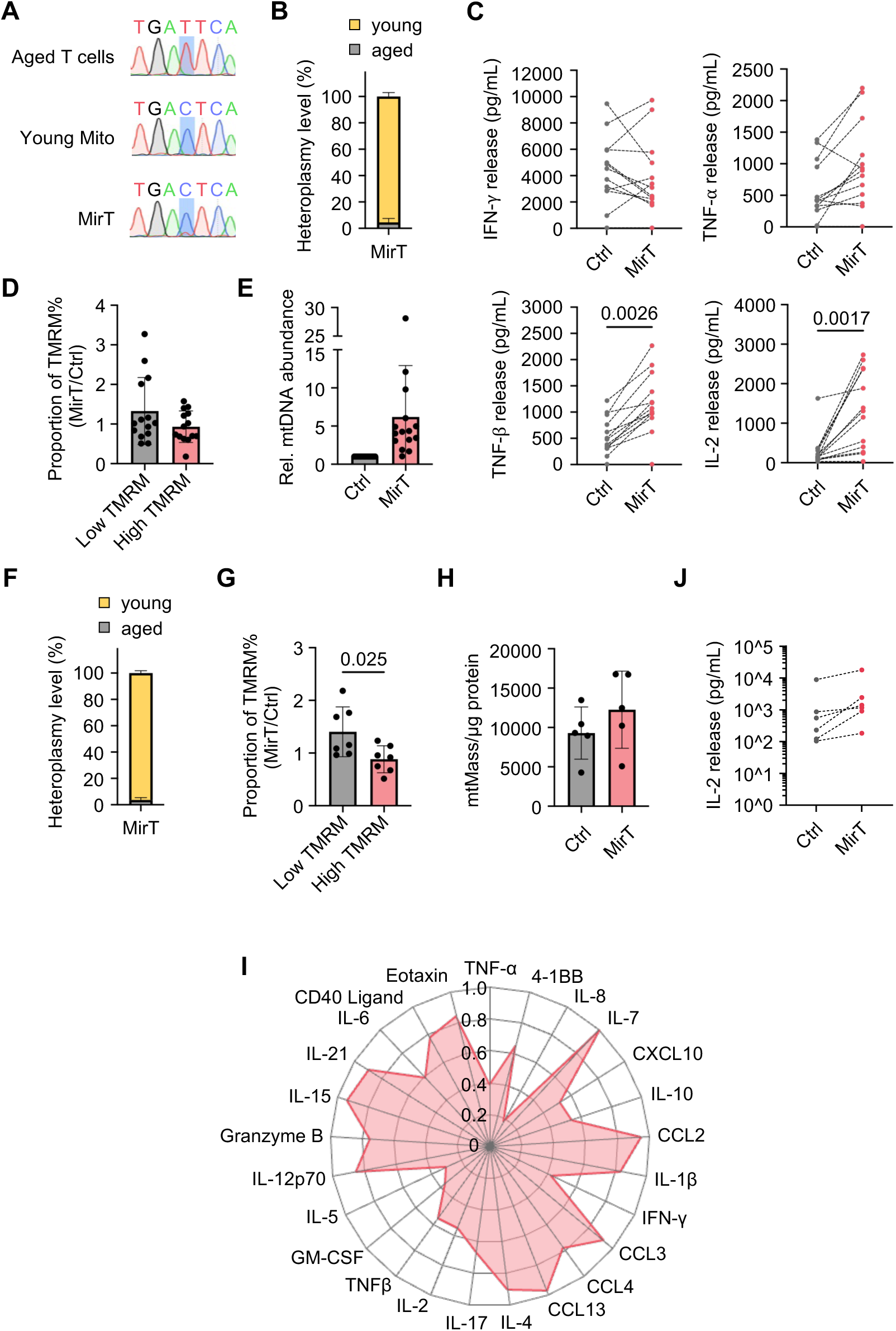
Functional restoration of aged human T cells following mitochondrial genome replacement. (A) Representative mtDNA sequencing chromatograms from aged human T cells, donor mitochondria from young cells, and mitochondrial genome-replaced T cells (MirT cells). (B) Estimated mtDNA heteroplasmy levels in MirT cells based on sequencing chromatogram signal intensities at donor- and recipient-specific mtDNA variants (n = 15). (C) Cytokine release following CD3/CD28 stimulation on day 7 after MirT cells generation (n = 13). (D) Relative proportions of low- and high-TMRM populations in MirT cells compared with control T cells, determined by flow cytometry (n = 14). (E) mtDNA content in MirT cells generated from T cells of elderly cancer patients, determined by quantification of the mitochondrial 12S rRNA gene (n = 9). (F) mtDNA heteroplasmy in MirT cells generated from T cells of elderly cancer patients, determined by mitochondrial genome sequencing (n = 9). (G) Relative proportions of low- and high-TMRM populations in MirT cells generated from T cells of elderly cancer patients (n = 7). (H) Mitochondrial mass in MirT cells generated from T cells of elderly cancer patients (n = 5). (I) Relative cytokine expression profiles in MirT cells generated from T cells of elderly cancer patients before stimulation, presented as MirT cells/Ctrl ratios (n = 8). (J) IL-2 release following CD3/CD28 stimulation in MirT cells generated from T cells of elderly cancer patients (n = 6). Data in (D), (E), (G), and (H) are presented as mean ± SD. Data in (J) are presented as median values. P values were determined using Student’s t-test.

### Mitochondrial genome replacement remodels transcriptomic programs in aged human T cells

To investigate transcriptomic alterations induced by mitochondrial genome replacement, bulk RNA-seq analysis was performed on untreated aged T cells (Ctrl) and MirT cells in cancer patients. Principal component analysis (PCA) demonstrated that Ctrl and MirT cells largely clustered together, indicating preservation of the overall transcriptional landscape following mitochondrial genome replacement (Fig. 7A). Nevertheless, differential expression analysis identified 23 upregulated and 82 downregulated genes in MirT cells compared with Ctrl cells (|log2FC| > 1, adjusted P value < 0.1) (Fig. 7B,C). Gene set enrichment analysis (GSEA) using the C7 ImmuneSigDB collection from MSigDB revealed enrichment of naïve- and resting T-cell-associated transcriptional signatures in MirT cells, including GSE11057_NAIVE_VS_EFF_MEMORY_CD4_TCELL_UP, GSE13738_RESTING_VS_TCR_ACTIVATED_CD4_TCELL_UP, and GSE26495_NAIVE_VS_PD1HIGH_CD8_TCELL_UP (Fig. 7D). In contrast, GSE36888_UNTREATED_VS_IL2_TREATED_TCELL_2H_DN was enriched in Ctrl cells. These findings suggest that mitochondrial genome replacement induces transcriptional remodeling toward less differentiated T-cell-associated programs. KEGG pathway enrichment analysis further demonstrated upregulation of pathways associated with protein synthesis and transcriptional regulation in eukaryotes (Fig. 7E). In parallel, pathways associated with fatty acid metabolism, including fatty acid metabolism and biosynthesis of unsaturated fatty acids, were enriched in MirT cells, whereas oxidative phosphorylation was reduced. In addition, inflammatory signaling-related pathways, including cytokine-cytokine receptor interaction and IL-17 signaling pathway, were relatively downregulated in MirT cells. Together, these findings indicate that mitochondrial genome replacement induces coordinated transcriptomic and metabolic remodeling in aged human T cells.

**Figure 7.**
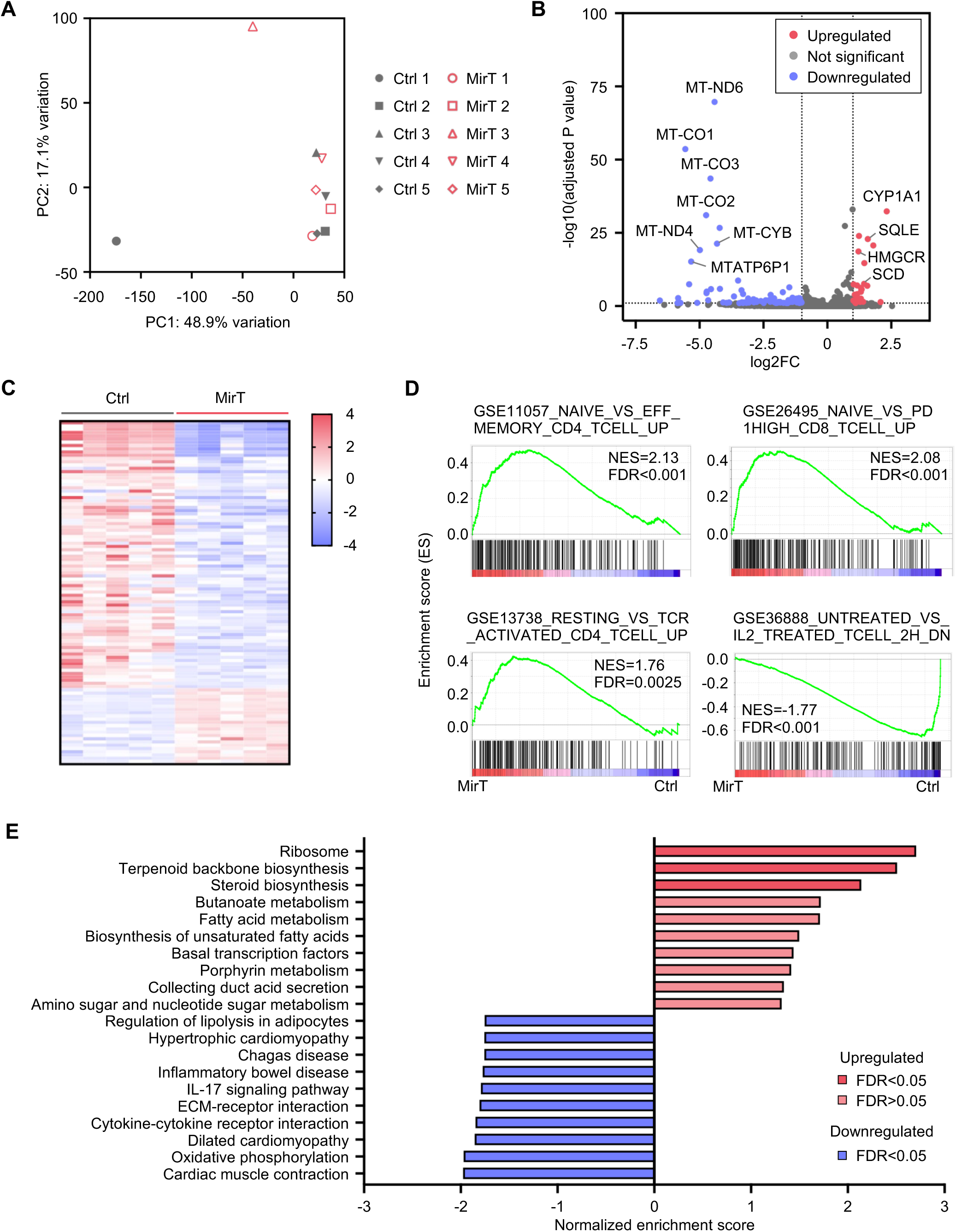
Transcriptomic remodeling toward youthful-like programs in mitochondrial genome-replaced aged human T cells. Bulk RNA-seq analysis of aged human T cells before and after mitochondrial genome replacement (MirT cells generation) was performed using paired samples from five donors. (A) Principal component analysis (PCA) showing transcriptomic distributions of control and MirT cell samples. (B) Volcano plot of differentially expressed genes (DEGs). Genes with log2 fold change > 1 or < −1 and adjusted P value < 0.1 were defined as DEGs. (C) Heat map showing hierarchical clustering of DEGs between control and MirT cell samples. (D) Gene set enrichment analysis (GSEA) using the ImmuneSigDB subset of the C7 collection from MSigDB, demonstrating enrichment of youthful and naïve T-cell-associated transcriptional programs in MirT cells. (E) KEGG pathway enrichment analysis using the complete ranked gene list.

## Discussion

In this study, we demonstrated in fibroblasts that mitochondrial genome replacement bidirectionally modulates aging-associated phenotypes. We next applied this approach to T cells, which play a central role in organismal aging, and found that replacement of aged mitochondrial genomes with young donor-derived mitochondrial genomes partially restored proliferative capacity and functional responsiveness, both of which are compromised during T-cell aging. Furthermore, adoptive transfer of mitochondrial genome-replaced T cells improved antitumor activity in tumor-bearing mice. In aged human T cells, mitochondrial genome replacement also induced transcriptomic remodeling toward youth-associated and less differentiated T-cell programs.

Mitochondria and the nucleus are linked through bidirectional signaling pathways that coordinate cellular metabolism, redox homeostasis, and gene expression (Grilo *et al*, 2021). Age-associated disruption of this mito-nuclear communication has been implicated in cellular dysfunction and loss of homeostasis (Zhu *et al*, 2022). Our findings raise the possibility that restoring mitochondrial genome integrity may partially re-establish mito-nuclear coordination and reverse aspects of cellular aging. Aging may not simply reflect stochastic damage accumulation, but rather progressive shifts in cellular aging trajectories propagated through interconnected metabolic, epigenetic, and inflammatory networks (Lopez-Otin *et al*., 2023). Mitochondrial dysfunction has been implicated both as a driver and amplifier of maladaptive signaling programs (Correia-Melo *et al*, 2016; Wiley *et al*, 2016). Notably, age-associated mitochondrial dysfunction cannot be fully explained by mtDNA mutations alone (Schon *et al*, 2012). Instead, mtDNA function depends on its organization within nucleoids, dynamic structures whose composition and regulation influence mitochondrial homeostasis (Bonekamp & Larsson, 2018; Kozhukhar & Alexeyev, 2023). We therefore propose that mitochondrial aging may be more appropriately understood in terms of “functional mitochondrial genetic integrity,” which reflects the integrated structural and regulatory state of the mitochondrial nucleoid system rather than DNA sequence fidelity alone. From this perspective, mitochondrial genome replacement may restore not only mtDNA sequence composition, but also broader layers of mitochondrial genetic regulation embedded within the nucleoid architecture. Although we did not directly measure functional mitochondrial integrity in the present study, mitochondrial genome replacement bidirectionally altered cellular aging phenotypes. Replacement of aged mitochondria with young mitochondria prolonged the time to growth arrest, whereas replacement of young mitochondria with aged mitochondria accelerated senescence-associated decline. Notably, epigenetic aging clocks are primarily based on nuclear molecular features (Hannum *et al*, 2013; Horvath, 2013). Therefore, the observation that mitochondrial replacement can modulate aging trajectories in both positive and negative directions further support the concept that mitochondrial functional state is tightly coupled to nuclear aging programs. These findings raise the possibility that functional mitochondrial state may act as a regulatory determinant of cellular aging dynamics. Recent studies have demonstrated that aging-associated phenotypes can be partially reversed through interventions targeting respiratory chain complex (Tavallaie *et al*, 2020), autophagy (Alsaleh *et al*, 2020), mitophagy (D’Acunzo *et al*, 2019), epigenetic remodeling (Ocampo *et al*, 2016), or cellular reprogramming pathways (Lu *et al*, 2020). In T cells, metabolic rejuvenation through enhancement of fatty acid oxidation has also been shown to partially restore antitumor function in aged cells (Al-Habsi *et al*, 2022). These observations collectively support the concept that cellular aging is not irreversibly fixed, but remains biologically plastic under certain conditions. Our findings extend this framework by suggesting that restoration of mitochondrial nucleoid integrity alone may be sufficient to bidirectionally influence aging-associated cellular states. Unlike nuclear reprogramming approaches that may compromise cellular identity, mitochondrial genome replacement may permit functional rejuvenation while preserving lineage-specific cellular programs.

Mitochondrial genomes exist in multiple copies within cells and form heterogeneous populations (heteroplasmy) at both intracellular and intercellular levels, which should be carefully distinguished when interpreting population-based analyses (Aryaman *et al*, 2018). To address this complexity, we previously established a single-cell ddPCR–based method capable of quantifying heteroplasmy at both levels (Maeda *et al*, 2020). In the present study, both murine and human MirT cells were predominantly composed of donor-genotype homoplasmic cells, suggesting that mitochondrial genome replacement may induce selective remodeling of mitochondrial genetic composition rather than passive coexistence of mixed mtDNA populations. This raises the possibility that selective pressures related to mito-nuclear compatibility and metabolic fitness influence post-replacement heteroplasmy dynamics (Lechuga-Vieco *et al*, 2020). An important unresolved issue is whether such remodeled mitochondrial states can be stably maintained long term. Although heteroplasmy remained stable for at least 16 days after replacement in our study, long-term persistence remains unknown. In reproductive mitochondrial replacement therapy for prevention of mitochondrial disease transmission in oocytes, reversion phenomena have been reported in which residual host mtDNA progressively re-expands despite initial donor predominance (Hudson *et al*, 2019), potentially through genetic drift or selective mito-nuclear interactions (Yamada *et al*, 2016). In contrast, our previous studies demonstrated stable maintenance of donor mitochondrial genotypes in iPS cells derived from mitochondrial genome–replaced fibroblasts (Maeda *et al*., 2021). Similar observations have been reported in pluripotent stem cells generated by somatic cell nuclear transfer (Ma *et al*, 2015). Collectively, these findings suggest that mitochondrial genome replacement is not merely passive mtDNA transfer, but a dynamic process of mitochondrial genetic integration into existing cellular systems.

Mitochondrial transfer and transplantation have attracted considerable attention as both physiological phenomena and therapeutic strategies (Brestoff *et al*, 2025; Dong *et al*, 2023). Clinical application of mitochondrial transplantation has raised concerns that isolated mitochondria exposed to extracellular calcium-rich environments may undergo structural disruption and release mitochondrial DAMPs (Giorgi *et al*, 2018; Hurst *et al*, 2017; Mills *et al*, 2017), potentially inducing inflammatory responses and limiting therapeutic safety (Chernyak, 2020). To minimize these risks, ex vivo approaches such as mitochondrial augmentation therapy were developed, in which isolated mitochondria are simply co-cultured with recipient cells prior to transplantation (Jacoby *et al*, 2022).

However, our findings suggest that simple addition of exogenous mitochondria may be insufficient for functional rejuvenation. In this study, simple mitochondrial co-culture induced detectable donor-derived heteroplasmy in T cells, yet failed to reproduce the proliferative restoration observed after mitochondrial genome replacement. Moreover, mitochondrial augmentation did not induce the enhanced mitophagy or mitochondrial biogenesis observed following MirT cells. These findings raise the possibility that transient mtDNA reduction followed by large-scale mitochondrial remodeling, rather than simple mitochondrial supplementation, may be required to reset aging-associated mitochondrial states. In this context, MirT cells may function as a mitochondrial remodeling trigger that promotes reorganization of mitochondrial genetic and metabolic networks.

Mitochondrial fitness and metabolic flexibility are essential for T-cell activation and function (Adamo *et al*, 2026; Han *et al*, 2023), yet how these properties change during aging remains incompletely understood (Luo *et al*, 2025; Mittelbrunn & Kroemer, 2021). Notably, studies of aged T cells have reported apparently conflicting metabolic phenotypes. For example, studies have reported increased spare respiratory capacity (Quinn *et al*, 2020), elevated basal respiration and ROS production (Yanes *et al*, 2019), or reduced OXPHOS and ATP production in aged T cells (Al-Habsi *et al*., 2022). Rather than representing mutually exclusive states, these observations may reflect distinct metabolic conditions shaped by T-cell subset, activation status, and environmental context. Collectively, these findings suggest that aging-associated T-cell dysfunction may reflect loss of metabolic coordination rather than simple energetic decline. In this context, elevated OCR may not necessarily reflect superior mitochondrial fitness, but may instead reflect maladaptive attempts to compensate for bioenergetic insufficiency. Our OCR analysis suggested that MirT cells did not simply enhance mitochondrial respiration. MirT cells normalized the elevated respiratory activity observed in aged T cells under homeostatic maintenance conditions toward a young-like range.

These findings suggest that functional rejuvenation may reflect restoration of metabolic coordination and flexibility rather than indiscriminate enhancement of OXPHOS. Importantly, MirT cells also exhibited improved cytokine production following stimulation, suggesting that normalization of basal metabolic activity did not impair inducible immune responsiveness. Rather, these findings raise the possibility that MirT cells restore a metabolically restrained yet functionally competent state characteristic of youthful T-cell populations. The transcriptomic changes observed after MirT cells further support the possibility that mitochondrial remodeling can secondarily reshape nuclear programs associated with T-cell aging and activation states. These findings suggest that the beneficial effects of MirT cells may arise not from simple enhancement of mitochondrial respiration, but from restoration of metabolic flexibility and relief from maladaptive hypermetabolic stress states associated with T-cell aging. Youthful T-cell function is defined not by maximal metabolic output, but by the ability to appropriately transition between quiescence and activation.

In the original MirT cells protocol using electroporation, aged murine T cells exhibited poor viability and limited cellular recovery following the procedure in line with previous reports describing reduced viability in engineered aged T-cell protocols (Kadyrzhanova *et al*, 2025). However, because murine aging models remain indispensable for mechanistic and long-term studies, we developed a simplified LNP-based MirT protocol that enabled efficient mitochondrial genome replacement in aged murine T cells while preserving both viability and cell number. This approach substantially simplified the preparation process compared with conventional LNP-based procedures (Li *et al*, 2025), reducing the need for extensive formulation and optimization steps. This protocol may facilitate future long-term studies investigating the persistence, safety, and biological effects of MirT cells. Several important questions remain unresolved. In the present study, we did not directly examine epigenomic remodeling downstream of mito-nuclear communication or the compatibility between nuclear and mitochondrial genomes after replacement. In addition, the mechanisms governing heteroplasmy shift, reversion, and stable maintenance of donor mitochondrial genomes, as well as the precise molecular alterations of mitochondrial nucleoids during aging, remain important subjects for future investigation. Collectively, our findings suggest that the mitochondrial genome represents not only a cellular unit profoundly altered during aging, but also a potential driver capable of bidirectionally influencing cellular fate. Replacement of aged mitochondria with young mitochondrial genomes therefore represents not merely a strategy to modify mitochondrial respiration, but rather an approach to restore metabolic coordination, flexibility, and cellular adaptability associated with youthful T-cell states.

## Materials and Methods

### Mice

C57BL/6J, C3H/HeNSlc, and NZB/NSlc mice were maintained under specific pathogen-free conditions. Experiments involving mouse breeding, maintenance, and tissue collection were performed at the Animal Experiment Center of Kyoto Prefectural University of Medicine. Adoptive T-cell transfer experiments using OT-1 mice were performed at Kyoto University. All animal procedures were approved by the respective institutional Animal Care and Use Committees of Kyoto Prefectural University of Medicine (Approval No. M2024-303) and Kyoto University (Approval No. Med Kyo 26353) and were conducted in accordance with institutional guidelines and regulations.

### Cells

CD3⁺ or CD8⁺ T cells were isolated from mouse spleens by negative selection using the EasySep™ Mouse T Cell Isolation Kit (STEMCELL Technologies, 19851) or EasySep™ Mouse CD8⁺ T Cell Isolation Kit (STEMCELL Technologies, 19853), respectively. Isolated T cells were cultured in RPMI 1640 medium containing L-glutamine (Nacalai Tesque, 30264-85) or TexMACS medium (Miltenyi Biotec, 130-097-196) at 37°C under 5% CO₂. Normal human dermal fibroblasts (NHDFs) were obtained from Lonza (CC-2511) or PromoCell (C-12302). Human fetal lung fibroblasts (TIG-1) and the human endometrial gland-derived mesenchymal cell line EPC100 were obtained from the Japanese Collection of Research Bioresources (JCRB) Cell Bank (JCRB0501 and JCRB1538, respectively). Cells were maintained in high-glucose DMEM (Fujifilm Wako Pure Chemical, 043-30085) supplemented with 10% fetal bovine serum (FBS) (Gibco, 10270-106) and 1% penicillin–streptomycin (Fujifilm Wako Pure Chemical, 168-23191) at 37°C under 5% CO₂.

### Human subjects

Human peripheral blood samples were obtained under protocols approved by the Institutional Review Boards of Kyoto Prefectural University of Medicine, Japan (Approval No. ERB-G-104-2), and Hadassah–Hebrew University Hospital (Approval No. 0296-22-HMO) and Galilee Medical Center (Approval No. 0202-22-NHR), Israel. Written informed consent was obtained from all participants prior to sample collection. Samples obtained at Kyoto Prefectural University of Medicine were used for the generation and validation of human MirT cells (Figure 3). Samples obtained at Hadassah–Hebrew University Medical Center were used for the characterization of MirT cells derived from aged healthy volunteers and aged cancer patients, including cytokine profiling and transcriptomic analyses (Figures 6 and 7).

### Plasmid DNA construction

The pCAGGS-MTS-XbaIR-Puro vector used in the fibroblast experiments contains the gene encoding the XbaI restriction endonuclease (GenBank: AF051092.1) and a puromycin resistance cassette downstream of the COX8A mitochondrial targeting sequence (Maeda *et al*., 2021). Plasmids for in vitro transcription were assembled using the NEBuilder HiFi DNA Assembly Master Mix (New England Biolabs, E2621S). pMRNAxp MTS-EGFP and pMRNAxp MTS-XbaI were generated by inserting a COX8-derived MTS and either EGFP or XbaI into the pMRNAxp mRNAExpress™ vector (System Biosciences, MR000PA-1) for in vitro transcription.

### mRNA synthesis

Plasmids were linearized with NdeI prior to transcription. In vitro transcription was performed using the mMESSAGE mMACHINE™ T7 ULTRA Transcription Kit (Invitrogen, AM1345) according to the manufacturer’s instructions. Reactions contained ATP (TriLink BioTechnologies, N-1501-25), CTP (TriLink BioTechnologies, N-1511-25), GTP (TriLink BioTechnologies, N-1512-25), N1-methylpseudouridine (TriLink BioTechnologies, N-1081-10), CleanCap Reagent M6 (TriLink BioTechnologies, N-7453-5), T7 reaction buffer, and T7 enzyme mix. Following incubation at 37°C for 2 h, template DNA was removed using TURBO DNase. Poly(A) tailing was subsequently performed using E-PAP enzyme according to the manufacturer’s protocol. Synthesized mRNA was purified using the Monarch RNA Cleanup Kit (New England Biolabs, T2040L) and stored at −80°C until use.

### Gene transfer

The pCAGGS-MTS-XbaIR-Puro vector was introduced into fibroblasts by electroporation using the Nucleofector Kit and Nucleofector 2b Device (Lonza, Walkersville, MD, USA). Two days after transfection, cells were subjected to puromycin selection (2 μg/mL) in DMEM containing 10% FBS and 1% penicillin–streptomycin for 24 h to enrich for transgene-expressing cells. T cells were transfected with in vitro-transcribed mRNA encoding the mitochondria-targeted XbaI restriction endonuclease using the ExPERT ATx electroporation platform (MaxCyte) with optimized electroporation programs according to the manufacturer’s instructions. Following electroporation, cells were immediately transferred to pre-warmed culture medium and incubated at 37°C under 5% CO₂. For LNP transfection, T cells were pre-activated for 1 day in TexMACS medium supplemented with recombinant human IL-7 (20 ng/mL; Miltenyi Biotec, 130-095-362) and CD3/CD28 Dynabeads™ (Gibco, 11453D). Magnetic beads were removed prior to transfection. Cells were then resuspended in TexMACS medium containing IL-7 (20 ng/mL) and recombinant human apolipoprotein E3 (ApoE3; FUJIFILM Wako Pure Chemical, 010-20261). Lipid nanoparticle (LNP)-formulated mRNA encoding the mitochondria-targeted XbaI restriction endonuclease (FUJIFILM) was added to the cultures and incubated overnight. Cells were subsequently maintained in TexMACS medium supplemented with IL-7, 10% FBS, and 1% penicillin–streptomycin. After delivery of mitochondria-targeted XbaI, cells were cultured in medium supplemented with uridine (50 μg/mL) until the initiation of mitochondria co-culture.

### Mitochondrial genome replacement protocol

Mitochondrial genome replacement was performed using a modified version of a previously reported protocol for fibroblasts (Maeda *et al*., 2021). Mitochondria were isolated from spleens obtained from C3H mice. Excised spleens were minced and transferred to gentleMACS C Tubes (Miltenyi Biotec, 130-093-237), followed by mechanical dissociation using the gentleMACS Dissociator program “m_mito_tissue_01”. Subtilisin A protease was added to the homogenate, which was incubated on ice for 10 min. Samples were centrifuged at 700 × g for 4 min, and the supernatant was collected. After addition of BSA, the suspension was sequentially filtered twice through a 40-μm filter and once through a 5-μm filter. Mitochondria were pelleted by centrifugation at 9,000 × g for 10 min and washed with respiration buffer containing 250 mM sucrose, 2 mM KH₂PO₄, 10 mM MgCl₂, 0.5 mM K-EGTA, and 20 mM K-HEPES.

Following the introduction of mitochondrial genome reduction constructs, fibroblasts or T cells were co-cultured with isolated mitochondria (40 μg mitochondrial protein per 1 × 10⁶ cells) at the appropriate time points for each cell type and gene-delivery method. Detailed timelines for each protocol are shown in Fig. 5B; EV1B, EV2A; and Appendix Fig. S1B. For T cells, cell–mitochondria mixtures were centrifuged at 1,500 × g for 5 min and resuspended in TexMACS medium supplemented with IL-7 (20 ng/mL), 10% FBS, and 1% penicillin–streptomycin. Fibroblasts were maintained under standard culture conditions, with medium replacement performed the following day. T cells generated by mitochondrial replacement were hereafter referred to as MirT cells.

### Evaluation of mitochondrial DNA copy number by SYBR Green qPCR

Mitochondrial DNA copy number (mtDNA copy number) was determined by quantitative PCR using a SYBR Green-based detection method. Genomic DNA was extracted using the NucleoSpin Tissue Kit (Macherey-Nagel), and 10 ng of DNA was used as template for each reaction. Real-time PCR was performed on a CFX Connect Real-Time PCR Detection System (Bio-Rad) using primer sets specific for the mitochondrial 12S rRNA gene and the nuclear β-actin gene (ACTB). Relative mtDNA copy number was calculated by normalizing the mitochondrial 12S rRNA signal to ACTB using the ΔΔCt method.

### Quantification of heteroplasmy by TaqMan SNP assay

Mitochondrial DNA (mtDNA) heteroplasmy was quantified using allele-specific TaqMan SNP assays. Real-time PCR was performed using 1 ng of genomic DNA and TaqMan™ Genotyping Master Mix (Applied Biosystems) on a CFX Connect Real-Time PCR Detection System (Bio-Rad). Allele-specific probes were designed to discriminate mtDNA variants of interest. Depending on the experimental system, informative mitochondrial polymorphisms between NHDF and TIG-1 cells, C57BL/6 and NZB/NSlc mice, human T cells and EPC100 cells, or C57BL/6 and C3H/HeNSlc mice were used to distinguish donor- and recipient-derived mtDNA molecules. mtDNA copy number was determined using standard curves generated from reference samples with known mtDNA copy numbers. Heteroplasmy levels were calculated as the proportion of donor-derived mtDNA relative to total mtDNA.

### Single-cell droplet digital PCR (sc-ddPCR) for microheteroplasmy

Single-cell mtDNA heteroplasmy was analyzed using droplet digital PCR (sc-ddPCR) as previously described (Maeda *et al*., 2020). Briefly, cells were loaded at a concentration previously validated to yield predominantly single-cell droplet occupancy, and approximately 500 cells per sample were analyzed following droplet generation using the QX200 Droplet Digital PCR System (Bio-Rad).

PCR amplification was performed using allele-specific primers and fluorescent probes targeting wild-type and mutant mtDNA sequences. Fluorescence signals were detected in two channels (FAM and VIC), and droplets were classified into four quadrants (double-negative, FAM-positive, VIC-positive, and double-positive) based on fluorescence thresholds established using control plasmids carrying the corresponding target sequences, as described previously (Maeda *et al*., 2020). Data were analyzed using QuantaSoft software (v1.7.4.917, Bio-Rad). Single-cell heteroplasmy was determined from the genotype classification of individual cell-containing droplets. Because the analysis was based on the presence or absence of donor and recipient mtDNA signals in individual cell-containing droplets rather than absolute molecule quantification, no Poisson correction was applied.

### Mitochondrial respiration analysis

Cellular respiration of fibroblasts was analyzed using an Oxygraph-2k high-resolution respirometer (Oroboros Instruments). Oligomycin (2 μg/mL), carbonyl cyanide p-trifluoromethoxyphenylhydrazone (FCCP; 1 μM), rotenone (0.5 μM), and antimycin A (2.5 μM) were sequentially added. Oxygen consumption rates were expressed as nmol O₂/mL and pmol O₂/s/10⁶ cells. Routine respiration, ATP-linked respiration, maximal respiration, spare respiratory capacity, proton leak, non-mitochondrial respiration, and coupling efficiency were calculated according to the manufacturer’s guidelines. Mitochondrial respiration of T cells was evaluated using an XFe96 Extracellular Flux Analyzer (Agilent Technologies). Cells were suspended in XF RPMI assay medium (Agilent Technologies, 103576-100) supplemented with 10 mM glucose, 1 mM pyruvate, and 2 mM L-glutamine and seeded at 1 × 10⁴ cells per well in Seahorse XFe96/XF Pro PDL Cell Culture Microplates (Agilent Technologies, 103799-100). Oligomycin (2 μM), FCCP (2 μM), and rotenone/antimycin A (0.5 μM) were sequentially injected using the Seahorse XF Cell Mito Stress Test Kit (Agilent Technologies, 103015-100). Oxygen consumption rate (OCR) was analyzed using Wave Controller software (v2.4, Agilent Technologies). Basal respiration, ATP production, maximal respiration, proton leak, spare respiratory capacity, non-mitochondrial respiration, and coupling efficiency were calculated according to the manufacturer’s instructions.

### RNA extraction, reverse transcription, and quantitative PCR

Total RNA was isolated using TRIzol reagent (Thermo Fisher Scientific, 15596018) followed by purification with the Direct-zol RNA MiniPrep Kit (Zymo Research, R2052) including on-column DNase I treatment. Reverse transcription was performed using 500 ng of total RNA and the PrimeScript RT Reagent Kit (Takara Bio, RR036A). Quantitative PCR was performed using KAPA SYBR Fast qPCR Master Mix (Kapa Biosystems, KK4602). Relative gene expression was normalized to Gapdh expression using the ΔΔCt method. Primer sequences are listed in Appendix Table S3.

### Adoptive T cell transfer experiment

CD8⁺ T cells were isolated from the spleens and lymph nodes of young (8–12 months old) or aged (16–17 months old) OT-1 transgenic mice. Cells were cultured for 5 days in vitro either for expansion or for MirT generation prior to adoptive transfer into aged C57BL/6 recipient mice. For all groups except the MirT group, recombinant mouse IL-7 (PeproTech, 217-17) was added at a final concentration of 50 ng/mL on days 1 and 3 of culture. On day 5, cells were harvested, counted, and adoptively transferred into recipient mice via tail vein injection at the indicated dose (5 × 10^5^, 5 × 10^4^, or 1 × 10^4^cells per mouse). MC38-OVA cells, an ovalbumin-expressing derivative of the MC38 colon adenocarcinoma cell line originating from C57BL/6 mice (H-2K^b^), were inoculated intradermally into the right flank of recipient mice on the same day as adoptive T-cell transfer. Anti-PD-L1 monoclonal antibody (clone 1-111A) was administered intraperitoneally at a dose of 25 μg per mouse every 6 days. Mice in the control group received intraperitoneal injections of PBS (200 μL per mouse) at the same schedule. Antitumor activity was evaluated by measuring tumor dimensions with calipers. Tumor volume was calculated using the following formula: Tumor volume (mm³) = 0.4 × length (mm) × width² (mm²).

### Clinical Study of Human MirT Cells and T-Cell Preparation

This non-interventional clinical study was conducted at Hadassah–Hebrew University Medical Center, Israel, to evaluate the feasibility and biological characteristics of MirT cells generated from T cells of elderly individuals using mitochondria isolated from peripheral blood mononuclear cells (PBMCs) of young healthy donors. Participants were enrolled into three groups: young healthy volunteers (18–35 years; n = 30), elderly healthy volunteers (>75 years; n = 9), and elderly patients with cancer (>65 years; n = 6). Eligible cancer patients had histopathologically confirmed solid tumors with evidence of advanced and/or metastatic disease. A single peripheral blood sample was collected from each participant. PBMCs obtained from young healthy volunteers were used as a source of donor mitochondria for mitochondrial replacement. CD3+ T cells isolated from PBMCs of elderly healthy volunteers and elderly cancer patients served as recipient cells for MirT generation and subsequent functional analyses. Clinical characteristics of the study participants are summarized in Appendix Table S1,S2. Recipient T cells were divided into two groups: one group underwent mitochondrial replacement to generate MirT cells, whereas the other was maintained as an untreated control (Ctrl). Control T cells were cultured at a density of 1 × 10^6^ cells/mL in RPMI 1640 medium (Sartorius, 01-100-1A) supplemented with MEM-α, 1× non-essential amino acids (NEAA; Gibco, 11140-035), and 50 μM 2-mercaptoethanol (Gibco, 31350-010) in a humidified atmosphere containing 5% CO2 at 37°C for 5 days. MirT cells and untreated control T cells were then cultured in triplicate in the absence or presence of ImmunoCult Human Anti-CD3/CD28 T Cell Activator (STEMCELL Technologies, 10971) at 3 μL/mL (1.5 μL per 0.5 mL culture) for 48 h prior to downstream cytokine and transcriptomic analyses.

### Cytokine Release Assay

Culture supernatants were collected after 48 h of culture with or without CD3/CD28 stimulation and subjected to cytokine analysis. Cytokine concentrations were measured using a Human Luminex Discovery Assay (25-Plex) panel (Bio-Techne R&D Systems, LXSAHM-25). Samples and standards were prepared and analyzed according to the manufacturer’s instructions, with up to 40 samples assayed per plate. Plates were read using a Bio-Plex MAGPIX multiplex reader system (Bio-Rad), and cytokine concentrations were calculated using the corresponding analysis software.

### RNA sequencing and analysis

Bulk RNA sequencing was performed on two experimental groups (n = 5 per group). Poly(A)-selected RNA libraries were prepared using the KAPA stranded mRNA-Seq Kit (Roche, KK8420) and sequenced as 122-bp single-end reads on an Illumina NovaSeq 6000 platform. Raw reads were quality-filtered and trimmed before alignment to the human reference genome GRCh38 (Ensembl release 106) using TopHat, allowing up to five mismatches. Gene-level counts were generated using htseq-count. Gene-level count matrices were generated from raw sequencing reads. Genes with fewer than 10 total counts across all samples were excluded. Filtered count data were imported into iDEP v2.4.4 for preprocessing and differential expression analysis (Ge *et al*, 2018). Low-abundance transcripts (<0.5 counts per million) were removed, and expression values were transformed using the EdgeR pipeline with a pseudocount of four. Differentially expressed genes were ranked according to log₂ fold change values. Gene Set Enrichment Analysis (GSEA; version 4.4.0, Broad Institute) was performed using a ranked list of all analyzed genes based on log₂ fold change values obtained from the differential expression analysis (Mootha *et al*, 2003; Subramanian *et al*, 2005).

Enrichment analysis was performed using the ImmuneSigDB (C7) collection from the Molecular Signatures Database (MSigDB) (Godec *et al*, 2016; Liberzon *et al*, 2015). Pathway enrichment analysis based on Kyoto Encyclopedia of Genes and Genomes (KEGG) (Kanehisa *et al*, 2021) annotations was additionally performed using WebGestalt 2024 (Elizarraras *et al*, 2024). RNA sequencing data have been deposited in the Gene Expression Omnibus (GEO) under accession number GSE336419.

### Senescence-associated β-galactosidase activity

Cells were incubated in phenol red-free HBSS containing SPiDER-βGal (1 μM; Dojindo, SG02) and DAPI (1 μg/mL; Dojindo, D523) at 37°C for 15 min. After washing, cells were resuspended in autoMACS Running Buffer (Miltenyi Biotec, 130-091-221) and analyzed by flow cytometry.

### Mitochondrial membrane potential and capacity]

Cells were stained with MitoTracker™ Deep Red FM (Invitrogen, M22426), tetramethylrhodamine methyl ester (TMRM; Invitrogen, I34361), and DAPI at 37°C for 20 min. Following washing, fluorescence was analyzed by flow cytometry.

### Mitochondrial reactive oxygen species

Cells were incubated with MitoSOX™ Red (Thermo Fisher Scientific, M36008) and DAPI in Hanks’ Balanced Salt Solution(-) without Phenol Red (Fujifilm Wako Pure Chemical, 085-09355) at 37°C for 20 min. Following washing, mitochondrial reactive oxygen species (mtROS) levels were quantified by flow cytometry.

### Assessment of T cell subsets and exhaustion markers

Cells were stained with fluorochrome-conjugated antibodies against CD44, CD62L, PD-1, TIM-3, and LAG-3 according to the manufacturers’ recommendations. Following incubation at 4°C for 30 min, cells were washed and analyzed by flow cytometry.

### Proliferation assay

Cell proliferation was assessed using the CellTrace™ Far Red Cell Proliferation Kit, for flow cytometry (Invitrogen, C34572). Labeled cells were cultured in TexMACS medium supplemented with IL-7 (20 ng/mL), 10% FBS, and 1% penicillin–streptomycin and stimulated with CD3/CD28 Dynabeads™ for 72 h. CellTrace fluorescence dilution was analyzed by flow cytometry and quantified using the Proliferation Platform in FlowJo version 10.10.0.

Division Index was defined as the total number of divisions divided by the number of cells at the start of culture. Proliferation Index was defined as the total number of divisions divided by the number of cells that went into division. Expansion Index was defined as the total number of cells divided by the number of cells at the start of culture. Replication Index was defined as the total number of divided cells divided by the number of cells that went into division.

### Statistical analysis

Data are presented as mean ± standard deviation (SD) unless otherwise indicated. Statistical analyses were performed using GraphPad Prism 11.0.2 (GraphPad Software). Comparisons among multiple groups were performed using one-way analysis of variance (ANOVA) followed by Tukey’s multiple-comparison test. Tumor growth curves were analyzed using two-way ANOVA followed by Tukey’s multiple-comparison test and are presented as mean ± standard error of the mean (SEM). P values < 0.05 were considered statistically significant.

## Ethics statement

This study was conducted in accordance with the Declaration of Helsinki and applicable institutional and national ethical guidelines. All human studies were approved by the respective Institutional Review Boards of the participating institutions.

## Data Availability

The RNA-sequencing data generated in this study have been deposited in the NCBI Gene Expression Omnibus (GEO) under accession number GSE336419. The data are currently private and will be made publicly available upon publication. During peer review, reviewers may access the data using accession number GSE336419 and the following token: azsvouuqlhsntib. Any additional information required to reanalyze the data reported in this paper is available from the lead contact upon request.

## Acknowledgements

We thank Kyoko Matsumoto and Atsushi Mizuno of IMEL Japan for fund accounting and administrative support, Yuki Kamiya for facilitating communication with the research facility in Israel, and Hiroshi Tamada for assistance with reporting to AMED. We are also grateful to Sayuri Shikata and Keiko Yurimoto for their technical assistance at Kyoto Prefectural University of Medicine and Kyoto University, respectively.

## Author contributions

Satoshi Gojo: Conceptualization; resources; supervision; validation; investigation; methodology; funding acquisition; writing–Original draft; writing- review and editing. Kenji Chamoto: Conceptualization; supervision; validation; investigation; methodology; writing–review and editing. Fumiya Teruyama: Conceptualization; formal analysis; investigation; methodology; writing–original draft; writing–review and editing. Toshihiko Taya: Investigation; methodology. Akira Shikuma: Investigation. Satoaki Matoba: Resources; funding acquisition. Kentaro Numajiri: Resources; Methodology. Sayako Umetani: Resources; Methodology. Yutong Chen: Investigation.

Amnon Peled: Resources; supervision; validation; investigation; methodology; funding acquisition. Eithan Galun: Investigation. Nimer Assy: Investigation. Aviad Zick: Investigation. Silvia Noiman: Resources; supervision. Dayana Michel: Resources; supervision. Dori Pelled: Resources; supervision; validation. Taro Inaba: Conceptualization; resources; supervision; Funding acquisition

## Expanded View Figure Legends

***Figure EV1.***

(A) Primer sets and probes used for the TaqMan SNP assay designed based on polymorphisms in the mtDNA D-loop region between NHDF and TIG-1 cells. Standard curves showing the amplification efficiency of the NHDF- and TIG-1-specific primer/probe sets are also shown.

(B) Experimental protocol for MirC generation. Recipient cells were transiently transfected with mitochondria-targeted XbaI (MTS-XbaI) to reduce endogenous mtDNA, followed by mitochondrial transfer and puromycin selection.

(C) Schematic illustration of aging induction and rejuvenation by mitochondrial genome replacement. Fibroblasts representing young or aged states were depleted of endogenous mitochondria and subsequently repopulated with mitochondria isolated from fibroblasts representing either young or aged states.

(D) Oxygen consumption rate (OCR) measured using the Oroboros O2k high-resolution respirometry system. Representative OCR traces of young fibroblasts and mock-transfected aged fibroblasts are shown. Blue line, oxygen concentration; red line, oxygen flux per cell. Oligomycin, FCCP, rotenone, and antimycin A were sequentially added at the indicated time points.

***Figure EV2.***

(A) Schematic illustration of the procedure used to generate mouse MirT cells.

(B) Representative chromatograms showing the mtDNA SNP (m.2766_2767AT>GC) distinguishing B6 and NZB mice.

(C) Primer sets and probes used for the TaqMan SNP assay designed based on the mtDNA SNP (m.2766_2767AT>GC) distinguishing B6 and NZB mice. The genomic location of the SNP and the positions of the primers and probes are indicated.

(D) Standard curves showing the amplification efficiency of the B6- and NZB-specific primer/probe sets used for mtDNA quantification.

***Figure EV3.***

(A) Representative mtDNA SNP chromatograms showing the sequence polymorphisms (m.16217T>C and m.16223C>T) distinguishing control T cells and EPC100 donor cells.

(B) Primer sets and probes used for the TaqMan SNP assay designed based on polymorphisms in the mtDNA D-loop region between control T cells and EPC100 cells. The positions of the SNPs, primers, and probes are indicated.

(C) Standard curves showing the amplification efficiency of the control T cell–and EPC100-specific primer/probe sets.

(D) Representative scatter plots of single-cell droplet digital PCR (sc-ddPCR) analysis of human MirT cells. Data shown are representative of N = 3 independent experiments.

***Figure EV4.***

(A) Representative flow cytometric analysis of EGFP expression in aged mouse CD3⁺ T cells following 0, 4, or 24 h of IL-7 exposure prior to transfection with MTS-EGFP mRNA. Histograms and the percentage of EGFP-positive cells at the indicated time points are shown.

(B) Experimental protocol for the generation of mitochondrial genome-replaced aged T cells (MirT cells). Unless otherwise indicated, MirT cells in subsequent figures refer to MirT cells generated by transfer of mitochondria from young donors into aged recipient T cells (MirT^Young/Aged^).

(C) Relative mtDNA copy number in aged mouse T cells before and 2 days after transfection with MTS-XbaI mRNA.

(D) Representative mtDNA SNP chromatograms showing the polymorphism distinguishing B6 and C3H mice.

(E) Primer sets and probes used for the TaqMan SNP assay designed based on the mtDNA polymorphism between B6 and C3H mice. The genomic location of the SNP (m.9348G>A) and the positions of the primers and probes are indicated. Standard curves showing the amplification efficiency of the B6- and C3H-specific primer/probe sets are also shown.

(F) Heteroplasmy levels in young and aged T cells and in MirT cells used for adoptive cell transfer (ACT) experiments.

